# The pre-existing human antibody repertoire to computationally optimized influenza H1 hemagglutinin vaccines

**DOI:** 10.1101/2021.10.25.465669

**Authors:** Kaito Nagashima, John V. Dzimianski, Julianna Han, Nada Abbadi, Aaron D. Gingerich, Fredejah Royer, Sara O’Rourke, Giuseppe A. Sautto, Ted M. Ross, Andrew B. Ward, Rebecca M. DuBois, Jarrod J. Mousa

## Abstract

The computationally optimized broadly reactive antigen (COBRA) approach has previously been used to generate hemagglutinin (HA) immunogens for several influenza subtypes that expand vaccine-elicited antibody breadth. As nearly all individuals have pre-existing immunity to influenza viruses, influenza-specific memory B cells will likely be recalled upon COBRA HA vaccination. We determined the epitope specificity and repertoire characteristics of pre-existing human B cells to H1 COBRA HA antigens. Cross-reactivity between wild type HA and H1 COBRA HA proteins were observed at both the oligoclonal B cell level and for a subset of isolated monoclonal antibodies (mAbs). The mAbs bound five distinct epitopes on the pandemic A/California/04/2009 head and stem domains, and the majority of the mAbs had HAI and neutralizing activity against pandemic H1 strains. Two head-directed mAbs, CA09-26 and CA09-45, had HAI and neutralizing activity against a pre-pandemic H1 strain. One mAb, P1-05, targets the stem region of H1 HA proteins, but does not compete with known stem-targeting H1 mAbs. We determined that mAb P1-05 recognizes a recently discovered membrane proximal epitope on HA, the anchor epitope, and we identified similar mAbs using B cell repertoire sequencing. In addition, the trimerization domain distance from HA was critical to recognition of this epitope by P1-05. Overall, these data indicate that seasonally vaccinated individuals possess a population of functional H1 COBRA HA-reactive B cells that target head, central stalk, and anchor epitopes, and demonstrate the importance of structure-based assessment of subunit protein vaccine candidates to ensure accessibility of optimal protein epitopes.

**Significance:** Influenza imposes significant human and economic costs every year. The current seasonal vaccine elicits primarily strain-specific antibodies, and year to year vaccine effectiveness is variable. The COBRA approach could provide longer protection and obviate the requirement for annual vaccination. Whereas COBRA HAs have previously been evaluated in animal models, the pre-existing COBRA HA-reactive human B cell population has yet to be elucidated, and is important to identify specific B cells that may be recalled by H1 HA COBRA vaccination. This work demonstrates that seasonally vaccinated individuals possess a functional B cell population targeting both head and stem domains that could be recalled with COBRA HA immunogens.

## Introduction

Influenza viruses are a major cause of morbidity and mortality worldwide each year (1). In particular, influenza A viruses (IAVs) and influenza B viruses cause annual epidemics in humans, and IAVs have caused multiple pandemics over the past century (2). Currently, H1N1 and H3N2 IAVs circulate in humans to cause epidemic disease (3, 4). Long-term protection to influenza viruses remains a challenge due to high mutation rates caused by a low-fidelity RNA polymerase, which leads to antigenic drift, as well as reassortment events of HA and NA with avian influenza viruses, which is termed antigenic shift (5). The mutability of influenza necessitates annual vaccination for protection against circulating strains (6). Current seasonal influenza vaccines provide protection against matched circulating viral strains. However, vaccine efficacy varies year to year due to mismatches between circulating strains and vaccine strains, as well as differences in hemagglutinin (HA) protein glycosylation patterns between vaccine and circulating strains (7–9). This variability in vaccine efficacy highlights the importance of developing an improved influenza vaccine, which would elicit an immune response to most circulating influenza A and/or B viruses (10). Current vaccines typically elicit strain-specific antibodies, and only a minority show cross-reactivity to other viral subtypes. The antibody response to influenza virus infection and vaccination focuses predominantly on HA. Within HA-targeting antibodies, those targeting the variable globular head domain dominate the response, whereas antibodies that bind the more conserved stem domain are elicited less frequently (11).

Computationally optimized broadly reactive antigen (COBRA) HA immunogens aim to elicit a broader antibody response compared to current seasonal vaccines (12, 13). In this approach, multiple-layered consensus building alignments of HA sequences are used to generate an immunogen encompassing multiple antigenic epitopes for one subtype (12). The resulting constellation of consensus epitopes, focused primarily in the antigenic sites of the head domain, represent diverse sequences that elicit broadly reactive antibodies in several animal models, including in mice and ferrets (12, 13). Structural analysis of COBRA HA immunogens has shown that these antigens resemble wild type HA proteins (14). The primary mechanism of COBRA HA-induced antibodies are through hemagglutination inhibition (HAI) and neutralization via HA head-domain binding antibodies (12). In contrast, stem-directed antibodies do not appear to be a major component of COBRA HA vaccine-induced immunity (15, 16).

H1N1 IAVs have caused two known pandemics, including the Spanish flu pandemic of 1918-1919, which caused an estimated 40-50 million deaths, and the 2009 swine flu pandemic, which caused an estimated 575,000 deaths (17). Circulating 2009 pandemic pH1N1/09-like viruses have replaced pre-2009 seasonal H1N1 influenza viruses in the human population (17). Antigenic sites defined on the H1 subtype HA have been characterized through mutagenesis studies in the presence of neutralizing antibodies (18). These highly variable sites are present on the immunodominant head domain, and include the Sa, Sb, Ca1, Ca2, and Cb sites (18). More recently discovered antibody epitopes include the receptor-binding site (RBS), the lateral patch, and the intratrimeric epitope, which exhibit broader reactivities (10, 19–21). A number of H1 subtype-based COBRA HAs have been previously described that incorporate both seasonal (pre- 2009) and pandemic-like (post-2009) influenza virus HA sequences. These include P1, which incorporates human sequences from 1933 to 1957 and 2009 to 2011 as well as swine sequences from 1931 to 1998, and X6, which incorporates human sequences from 1999 to 2012 (22).

An individual’s immune history to influenza also plays a major role in the antibody response to vaccination. For example, the idea of original antigenic sin (OAS) describes the dominant nature of the antibody response to the first influenza virus strain compared to exposures to subsequent strains (23). While COBRA HA immunizations have been shown to be efficacious in naïve as well as pre-immune mouse and ferret models of influenza infection, pre-existing immunity to COBRA HAs in humans has not been investigated, and is important to understand as these antigens move toward clinical trials. Here, we identify epitope and repertoire characteristics of the recalled antibody response from previous infection and vaccination that may be stimulated by H1 COBRA HA antigens. We show that human antibodies and B cells that cross-react with COBRA HAs are present in individuals vaccinated with the 2017-2018 and 2019-2020 quadrivalent influenza vaccine (QIV). A panel of 26 monoclonal antibodies (mAbs) was isolated, and these mAbs bind five distinct epitopes on the A/California/04/2009 HA protein, including an epitope near the viral membrane, termed the anchor epitope. Moreover, a subset of these mAbs bind both pre- and post-2009 pandemic strains with demonstrable HAI and neutralization activity. Overall, our data identify the major epitopes and repertoire characteristics of pre-existing human antibodies that recognize COBRA HA antigens.

## Materials and Methods

### Human subject samples

All human studies were approved by the University of Georgia Institutional Review Board. mAb isolation was conducted from subjects vaccinated with the 2017-2018 seasonal influenza vaccine (Fluzone) from peripheral blood mononuclear cells (PBMCs) isolated from blood draws 21-28 days following vaccination. Repertoire sequencing was completed from a single human subject vaccinated with the 2019-2020 influenza vaccine (Fluzone) from blood obtained 28 days following vaccination.

### B cell expansion of human subject PBMCs

PBMCs were plated at a density of 25,000 cells/well in a 96-well plate on a layer of gamma-irradiated NIH 3T3 cells (20,000 cells/well) expressing hCD40L, hIL-21, and hBAFF in the presence of CpG and cyclosporine A as previously described (24, 25). B cell supernatants were screened by enzyme-linked immunosorbent assay (ELISA) at 7 days post-plating of PBMCs.

### Expression and purification of recombinant influenza HA proteins

Trimeric wild-type HA or COBRA HA ectodomains were expressed and purified in Expi293F cells following the manufacturer’s protocol and as previously described (26). Collected supernatants containing the HA antigens were purified on a HisTrap Excel column following the manufacturer’s recommended protocol. Eluted fractions were pooled and purified proteins were verified for integrity by probing with an anti-HIS tag antibody (Biolegend) as well as with subtype-specific mAbs via SDS-PAGE and Western blot.

### ELISA screening of B cells, hybridoma supernatants, and mAbs

Untreated 384-well plates (VWR) were coated with recombinant HA proteins diluted to 2 μg/mL in PBS at 4 °C overnight. Plates were washed once with water, then blocked with 2% blocking buffer (PBS + 2% non-fat dry milk (Bio-Rad) + 2% goat serum + 0.05% Tween-20) for 1 hr at room temperature. Plates were washed three times with water, and 25 µL of B cell supernatants, hybridoma supernatants, or mAbs were added. mAbs were serially diluted three-fold in PBS from 20 μg/mL prior to addition for twelve total dilutions. Plates were incubated at 37 °C for 1 hr, then washed three times with water. Goat anti-human IgG Fc-AP secondary antibody (Southern Biotech), diluted 1:4000 in 1% blocking buffer (1:1 dilution of PBS and 2% blocking buffer), was added and plates were incubated at room temperature for 1 hr. Plates were then washed five times with PBS-T (PBS + 0.05% Tween-20). *p*-Nitrophenyl phosphate (PNPP) substrate, diluted in substrate buffer (1.0 M Tris + 0.5 mM MgCl_2_, pH=9.8) to 1 mg/mL, was added, and plates were incubated for 1 hr and read at 405 nm on a BioTek plate reader. To quantify HA-reactive IgG from each subject, plates were coated overnight with eight two-fold serial dilutions of human plasma IgG standard (Athens Biotechnology) starting at 10 μg/mL. All steps were followed as for antigen, except PBS was used in the primary antibody step. GraphPad Prism was used to interpolate antigen-reactive IgGs from the human plasma IgG standard curve. The EC_50_ value for each mAb was determined by using the four-parameter logistic curve fitting function in GraphPad Prism software.

### Generation of HA-reactive mAbs

Eight days following plating of PBMCs, wells identified to contain positive B cells by ELISA were selected for electrofusion to generate hybridomas as previously described (24, 25). Hybridomas were plated in 384-well plates for HAT selection, and grown for 14 days at 37°C, 5% CO_2_. Following screening by ELISA, hybridomas were single-cell sorted using a MoFlo Astrios cell sorter using live/dead staining by propidium iodide. The sorted hybridomas were cultured in 25% Media E (StemCell) + 75% Media A (StemCell) for two weeks, then subjected to another round of screening by ELISA. Hybridomas with the highest signal were grown in 250 mL serum-free media (Gibco) for approximately one month. Secreted mAbs were purified using a Protein G column (GE Healthcare) and concentrated for use in downstream assays.

### Hybridoma sequencing

Hybridoma cell lines encoding each mAb were sequenced utilizing the primers described by Guthmiller *et al.* (27). Briefly, RNA was extracted from each hybridoma and cDNA was generated using the SuperScript IV First-Strand cDNA Synthesis Kit (Invitrogen). A nested PCR protocol was used to generate sequencing products. In the first nested PCR step, a primer mix specific to the heavy, kappa, or lambda chain *V* gene and the constant region were used to amplify the variable region using the cDNA as template. In the second PCR step, the first PCR product was used as a template with a nested primer mix to improve product specificity and yield. The second nested PCR products were sequenced using the constant region primer and the *V, D,* and *J* alleles were identified by IMGT/V-QUEST (28).

### Hemagglutination inhibition assay

The HAI titer for each mAb was determined as previously described (16). Influenza viruses were titered to eight HAUs (hemagglutination units). 50 μL of mAbs diluted to 20 μg/mL in PBS were added to the first well of a 96-well U-bottom plate (VWR), and diluted two-fold in PBS for 25 μL mAb total per dilution. Eight HAUs of virus were added in a 1:1 ratio to each mAb dilution, and each well was mixed and incubated for 20 min at room temperature. Following this, 50 μL of 1.0% turkey red blood cells (Lampire) were added per well. Plates were read 45 min after the addition of 1.0% turkey red blood cells.

### Focal reduction assay

Focal reduction assays (FRAs) were completed for each mAb as previously described (16). MDCK cells were plated in 96-well plates overnight to achieve >95% confluency the next day. Cells were washed twice with PBS, and 50 µL of virus growth media (VGM: DMEM + 2 μg/mL TPCK-trypsin + 7.5% BSA) were added and the plates were returned to the incubator at 37°C, 5% CO_2_. mAbs at 20, 8, or 1 μg/mL were serially diluted two-fold in VGM, and virus was diluted to a concentration of 1.2x10^4^ FFU/mL in VGM. MDCK cells were washed with PBS and 25 μL serially diluted mAbs were added, followed by 25 μL of 1.2x10^4^ FFU/mL of virus. Plates were incubated at 37 °C, 5% CO_2_ for 2 hr, and then 100 μL/well of overlay media (1.2% Avicel + modified Eagle media (MEM)) were added and incubated overnight. The overlay was removed and wells were washed twice with PBS. Ice-cold fixative (20% formaldehyde + 80% methanol) was added and plates were incubated at 4 °C for 30 min. Plates were washed twice with PBS and permeabilization buffer (PBS + 0.15% glycine + 0.5% Triton-X 100) was added, followed by a 30 min incubation. Plates were washed three times with PBS-T and primary IAV anti-NP mouse antibody (IRR), diluted 1:2000 in ELISA buffer (PBS + 10% goat serum + 0.1% Tween-20), was added. Plates were incubated at room temperature for 1 hr. Plates were then washed three times with PBS-T and secondary goat anti-mouse IgG-HRP antibody (Southern Biotech), diluted 1:4000 in ELISA buffer, was added. Plates were incubated at room temperature for 1 hr and then washed with PBS-T. KPL TrueBlue Peroxidase substrate was added per well and plates were incubated for 10-20 min. Plates were washed, dried, and foci were enumerated using an ImmunoSpot S6 ULTIMATE reader with ImmunoSpot 7.0.28.5 software (Cellular Technology Limited). Neutralizing IC_50_s were calculated using the GraphPad Prism four-parameter logistic curve fitting function.

### Epitope binning by biolayer interferometry

The panel of mAbs isolated from human subjects were competed for binding using the A/California/04/2009 HA protein on the OctetRED384 system as previously described (24). Anti-penta-HIS biosensors (Sartorius) were immersed in kinetics buffer (PBS + 0.5% BSA + 0.05% Tween-20) for 60 s to obtain a baseline reading. Biosensors were then loaded with 100 μg/mL of A/California/04/2009 HA protein diluted in kinetics buffer for 60 secs. Biosensors were returned to kinetics buffer for a baseline of 60 s. Following this, biosensors were immersed in the first mAb (100 μg/mL in kinetics buffer) for 300 s for the association step. The biosensors were then immersed in the competing, second mAb (100 μg/mL in kinetics buffer) for 300 s. The biosensors were then regenerated in 0.1 M glycine, pH = 2.7 and PBS alternately for three cycles before proceeding to the next mAb competition set. The extent of competition was calculated as the percentage of the signal from the second mAb in the second association step in the presence of the first mAb to that of the second mAb alone in the first association step for all biosensors. A ratio of <=33% was considered complete competition, >33 and <=67% moderate competition, and >67% no competition.

### Antibody escape mutant generation

To identify epitopes important for mAb binding, MDCK cells were plated in 24-well plates overnight to achieve >95% confluency. Cells were washed twice with PBS, then 200 μL of virus, diluted in VGM to an MOI=0.01, was added. Cells were returned to 37 °C, 5% CO_2_. Infection proceeded for 1 hr. Viral inoculum was then removed and VGM containing mAb at 1x IC_50_ was added. Cells were returned to 37 °C, 5% CO_2_ for 24 hr. For the subsequent passage, MDCK cells at >95% confluency were washed with PBS twice, and a 1:10 dilution of the media from the first passage in 200 μL virus growth media was used to infect MDCK cells, followed by a 1 hr incubation. Viral inoculum was removed and virus growth media containing mAb at 1x IC_50_ was added, and cells were incubated at 24 hr at 37 °C, 5% CO_2_. This was done for five passages with mAb at 1x IC_50_, then one passage with mAb at 2x IC_50_, followed by two passages with mAb at 3x IC_50_ for eight passages total. After the eighth passage, media containing escape mutant virus was stored, and cells were pelleted by centrifugation at 3000 rpm. Cells were lysed using SPRI beads and RNA was extracted. vRNA encoding the HA gene was amplified using the Uni12 primer specific to all influenza genome segments by RT-PCR (29). In a subsequent PCR step, HA-specific primers from Deng et al. (29) were used to amplify the HA segment of the pandemic H1N1 virus. Sanger sequencing was used to identify escape mutations. Escape mutations were considered significant either if they appeared in two of three replicates, or if a mutation was present within ten amino acids of another.

### Antibody-dependent phagocytic activity of mAbs

To measure antibody-dependent phagocytic activity, 2×10^9^ 1-µm Neutravidin-coated yellow-green FluoSpheres (Invitrogen #F8776) were resuspended in 1 mL of 0.1% PBS. The FluoSpheres were then centrifuged at 5000 rpm for 15 minutes, 900 μL supernatant was removed, and the FluoSpheres were resuspended with 900 µL of 0.1% PBS. This process was repeated for a second wash, then the FluoSpheres were resuspended with 20 µg of biotinylated Y2 protein. The FluoSpheres were then incubated overnight at 4 °C, protected from light, with end-to-end rocking. Next, HA-specific antibodies were diluted in complete RPMI media (cRPMI, RPMI + 10% FBS) to a final concentration of 1 µg/mL in a U-bottom 96-well plate. Then, 20 µL of antibody dilution was transferred into a clean F-bottom 96-well plate, and 10 µL of FluoSpheres were added with the antibody followed by a 2 hr incubation at 37 °C for opsonization. After 1.5 hr, THP-1 cells were centrifuged at 200 × g for 5 min, washed once with PBS, then resuspended in culture medium (RPMI & 10% FBS) at a concentration of 5×10^5^ cells/mL. Then, 200 µL of cells were added to each well and incubated at 37 °C with 5% CO_2_ while shaking for 6 hr. Once the incubation finished, the plate was then centrifuged at 2000 rpm for 5 min. Then, 100 µL was pipetted out of each well and replaced with 100 µL of cold 4% paraformaldehyde to fix the cells. The plate was then left at room temperature for 20 min, protected from light. The plate was then stored at 4 °C in the dark. Cells were then analyzed with a NovoCyte Quanteon flow cytometer.

### Expression and purification of recombinant proteins for electron microscopy

For EM studies, Y2 HA COBRA was cloned using Gibson assembly into a derivative of pcDNA3.1+ (30). Plasmids for the P1-05 heavy and light chain were synthesized (Genscript) and cloned into pcDNA3.1+. Cells and media were purchased from Thermo Fisher Life Technologies unless stated otherwise. Y2 HA protein expression was initiated by transfection of endotoxin free DNA into CHO-S cells using flow electroporation technology (MaxCyte). Transfected cells were suspended in CD OptiCHO supplemented with 2 mM GlutaMAX, HT, 0.1% pluronic acid, and incubated at 37 °C, 8% CO_2_, 85% humidity in an orbital shaker (Kuhner). After 24 hrs, cultures were supplemented with 1 mM sodium butyrate, and the culture temperature was dropped to 32°C. Cultures were supplemented daily with MaxCyte CHO A Feed (0.5% yeastolate, 2.5% CHO-CD Efficient Feed A, 2 g/L glucose, 0.25 mM GlutaMAX). The media was harvested 8-12 days post-transfection and filtered. For purification of Y2, media was diluted with an equal volume of Buffer A (500 mM NaCl, 20 mM sodium NaH_2_PO_4_, 20 mM imidazole) and loaded onto a 1 mL HisTrap column (GE Healthcare). The column was washed with Buffer A and the protein eluted with a gradient to Buffer B (500 mM NaCl, 20 mM NaH_2_PO_4_, 500 mM imidazole) on an ÄKTA Pure chromatography system (GE Healthcare). Fractions containing the protein were pooled, concentrated, and further purified and buffer exchanged on a Superdex 200 10/300 column (GE Healthcare) equilibrated in PBS (Sigma). Fractions were pooled and concentrated, then flash frozen in liquid N_2_ and stored at -80°C until use. For P1-05, the mAb was purified using a 1 mL HiTrap Protein A HP column (GE Healthcare). The media was diluted with an equal volume of Protein A IgG Binding Buffer (Thermo Scientific) and loaded onto the column. The column was washed with binding buffer, then eluted with a gradient to Protein A IgG Elution Buffer (Thermo Scientific). To adjust the pH, 55 μl of 1.89 M Tris pH 8 was added per 1 mL fraction. Fab was generated and purified using the Pierce™ Fab Preparation Kit according to the manufacturer’s instructions (Thermo Scientific). The Fab product in PBS was flash frozen in liquid N_2_ and stored at -80 °C until use.

### Cloning and expression of Y2 COBRA with a thrombin cleavage site

Y2 COBRA was cloned into the pBacPAK8 vector in frame with an N-terminal gp67 signal sequence and C-terminal thrombin cleavage site, T4 fibritin domain, and hexahistidine/StrepTag II tags. The construct design results in predicted vector supplied sequences of AATNA and LVPRGSPGSGYIPEAPRDGQAYVRKDGEWVLLSTFLGHHHHHHGGSWSHPQFEK at the N-and C-termini, respectively. Baculovirus was generated using the *flash*Bac™ kit according to the manufacturer’s instructions (Mirus Bio). The protein was expressed in 2 L of Sf9 cells at 2×10^6^ cells/mL maintained in ESF921 media (Expression Systems) by adding 25 mL virus per liter of culture. The media was harvested after 3 days, pH adjusted with NaCl and Tris pH 8, and stored at -20°C. Prior to purification, the thawed media was filtered and concentrated to 150-200 mL by tangential flow with a Vivaflow® 200 (Sartorius). The resulting sample was diluted with an equal volume of Buffer A, filtered, and loaded onto a 5 mL HisTrap column (GE Healthcare). The column was washed with Buffer A and the protein eluted with a gradient to Buffer B. The protein was pooled, concentrated, and supplemented with 5% glycerol prior to flash freezing and storage at - 80 °C.

### Kinetic assays by biolayer interferometry

Biolayer interferometry kinetic assays were performed in triplicate on the Octet® Red384 system (Sartorius) with a buffer containing PBS, 1% BSA, and 0.05% Tween. Anti-penta-HIS biosensors were immersed in buffer for 120 s, then loaded with 10 μg/mL Y2 for 300 s. The biosensors were then dipped into buffer for 120 s to obtain a baseline, dipped into buffer containing P1-05 Fab in a dilution series ranging from 54 nM to 0.67 nM for 300 s, and buffer for 600 s to measure dissociation. The data were processed in the Octet Data Analysis HT software v7 (Sartorius). Each curve was reference subtracted, aligned to the baseline, and aligned for inter-step correction through the dissociation step for each curve. Each replicate was fit globally for well-resolved curves in the dilution series using a 1:1 binding model. Parameters were optimized based on the R^2^, χ^2^, and individual *K*_D_ error values to maximize the goodness of fit. The final reported *K*_D_ value (98.5±32.3 pM) represents the mean ± standard deviation of three independent experiments.

### Electron microscopy of the Y2+P1-05 complex

The protein samples were thawed on ice. To generate the immune complex, P1-05 Fab and Y2 COBRA produced in CHO cells were combined in a 3:1 Fab:HA trimer ratio and incubated at room temperature for 1 hr. For negative stain analysis, the immune complex was deposited at 15 μg/mL onto carbon-coated, glow-discharged, 400 mesh copper grids (EMS) and stained with 2% w/v uranyl formate. The sample was imaged on an Arctica Talos 200C electron microscope (FEI) operating at 73,000x nominal magnification with a Falcon II direct electron detector and a CETA 4k camera (FEI). Micrographs were collected with Leginon and particles were picked using a difference of Gaussians particle picker and processed with Appion (31–33). Particles were classified in 2D and 3D in Relion 3.0 and Cryosparc2 and reconstructed in 3D in Cryosparc2 (34, 35). Figures were made in UCSF Chimera (36).

### Single-cell V(D)J sequencing and analysis

PBMCs were stained with the following antibodies and proteins for flow sorting: anti-CD19-APC (1:10 dilution, clone HIB19, cat. no. 982406, BioLegend), anti-IgD-FITC (1:20 dilution, clone IA6-2, cat. no. 348206, BioLegend), anti-IgM-FITC (1:20 dilution, clone MHM-88, cat. no. 314506, BioLegend), Ghost Dye Red (1:1000), Y2-PE (1:20 dilution), and Y2-BV605 (1:20). AviTagged Y2 COBRA HA proteins containing the Y98F mutation to reduce sialic acid binding were biotinylated using the BirA biotin-protein ligase in the BirA500 kit (Avidity) and complexed to streptavidin-fluorophores SA-PE (1:500 dilution, cat. no. S866, Thermo Fisher) and SA-BV605 (1:250 dilution, cat. no. 405229, BioLegend). CD19^+^IgM/IgD^-^PE^+^BV605^+^ double-positive, antigen-specific B cells were flow sorted on the MoFlo Astrios and resuspended in PBS+0.04% BSA. These cells were then used to generate Single Cell 5’ v2 Dual Index V(D)J libraries using the 10X Chromium Next GEM Single Cell 5’ Reagent Kit v2 (10X Genomics). Libraries were then sequenced using a NextSeq 550 sequencer (Illumina). Single-cell V(D)J FASTQ files were generated and demultiplexed using Cell Ranger v4.0.0 and data were visualized using the Loupe VDJ v3.0.0 browser. Only B cells with intact variable regions and paired heavy and light chains were considered for downstream analysis.

## Results

### COBRA HA-specific B cell responses

To determine the size of the H1 COBRA HA-reactive B cell population within seasonally vaccinated individuals, total B cells from four vaccinated subjects (2017-2018 cohort) were stimulated on an irradiated feeder layer as previously described (24). B cell supernatants were assayed for activity against A/California/04/2009 HA, P1 COBRA HA, and X6 COBRA HA recombinant proteins by ELISA. As expected, HA- and COBRA HA-reactive IgG titers were higher in day 21 post-vaccination samples compared to those obtained pre-vaccination **(Figure 1).** Comparisons of A/California/04/2009 HA-reactive IgG titers to those of P1 HA- and X6 HA-reactive IgGs indicated that binding to A/California/04/2009 HA protein was consistently higher. The majority of subjects demonstrated significant P1 HA-reactive IgG titers that, while lower than A/California/04/2009 HA-reactive IgG titers, were higher than or equivalent to X6 HA-reactive IgG titers in three of four subjects. The disparity in antibody titers between natural A/California/04/2009 HA- and COBRA HA-reactive proteins may be attributed to the relatively high abundance of pandemic strain-specific antibodies, and the absence of these potential binding epitopes on the H1 COBRA HA proteins. Moreover, the degree of similarity of each COBRA HA to the A/California/04/2009 HA appeared to be reflected in the degree to which reactive IgG titers were elicited. Namely, P1 COBRA HA, representing pandemic-like human and swine H1 HA sequences, demonstrates 84.63% identity to A/California/04/2009 HA, whereas X6 HA, representing seasonal-like H1 HA sequences, demonstrated a lower 80.53% identity to A/California/04/2009 HA.

**Figure 1.**
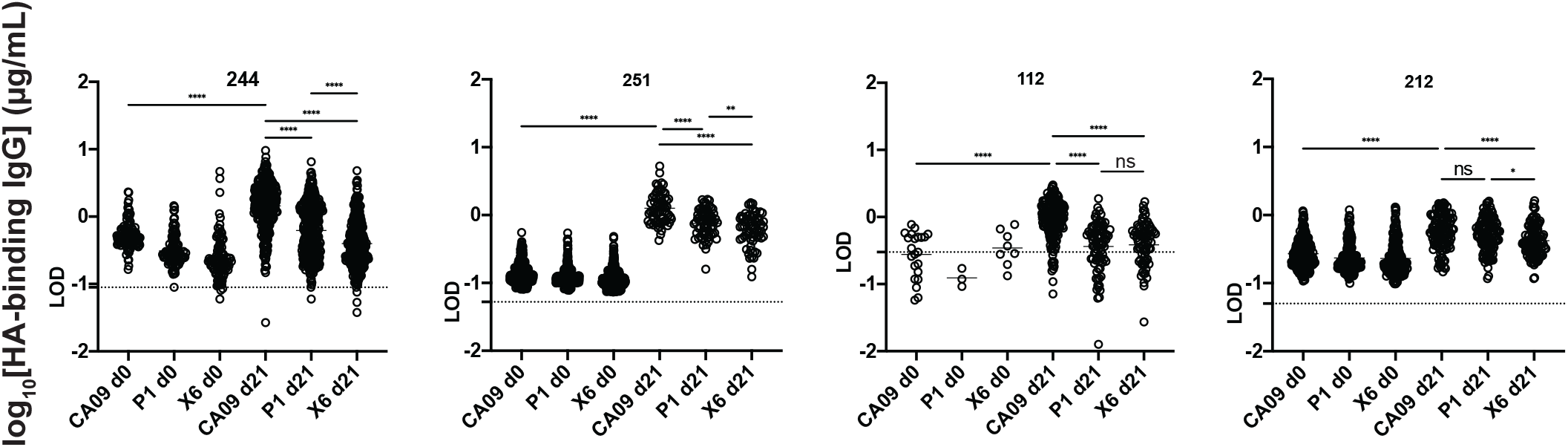
Binding titers of oligoclonal B cell supernatants pre-vaccination (d0) and 21 days post-vaccination (d21) from four representative subjects. IgG titers against CA09 HA (CA09), P1 COBRA (P1), and X6 COBRA (X6), are shown for representative subjects receiving the 2017-2018 QIV. Supernatants from stimulated PBMCs were screened by ELISA using plates coated with the indicated antigen. PBMCs were standardized to 25,000 cells per well. Each circle indicates 1 well, the mean is shown as a bar, and the limit of detection (LOD) is indicated by a dotted line. LOD was calculated as three times the standard deviation of the lowest concentration divided by the slope of the standard curve for each sample, interpolated to its corresponding log concentration. ****P<0.0001, **P=0.0062, *P=0.0216, ns=not significant.

### Lineage analysis of pre-existing COBRA HA-specific mAbs

To further probe the pre-existing B cell response to COBRA HA antigens, we isolated 26 mAbs from five additional human subjects vaccinated with the 2017-2018 quadrivalent influenza vaccine within the same cohort using A/California/04/2009 HA and P1 COBRA HA as screening antigens. The antibody-encoding genes were sequenced, and the results indicated the usage of a diverse set of immunoglobulin *V* genes across the entire panel **(Figure 2, Table S1)**. When comparing usage of heavy chain genes, *V*_H_1, *V*_H_3, *V*_H_4, and *V*_H_5 gene families were represented **(Figure 2A)**. Approximately 50% of all mAbs utilized a gene from the *V*_H_*3* family, and approximately another 50% utilized a gene from the *V*_H_*4* family. In the light chain, for those mAbs utilizing the kappa chain, a significant proportion utilized genes *V*_K_*3-11* and *V*_K_*3-15*. The remainder used *V* genes *V*_K_*3-20* or those from *V_K_1* or *V_K_2* families. mAbs utilizing the lambda chain used predominantly *V*_L_*3-21* and *V*_L_*2-14*. Paired heavy and light chain *V* genes showed variation across the antibody panel, with the *V*_H_*3-7*:*V*_K_*3-15* and *V*_H_*4-39*:*V*_L_*2-14* pairings being the most abundant for kappa- and lambda chain-utilizing mAbs, respectively **(Figure 2B)**. The lengths of the heavy and light chain junctions ranged from 12-24 amino acids for the heavy chain, 10-12 amino acids for the kappa chain, and 12-14 amino acids for the lambda chain **(Figure 2C).** The percent identities of the variable genes to the germline sequence had averages of 93% for both the heavy and kappa chains, and 96% for the lambda chain **(Figure 2C)**.

**Figure 2.**
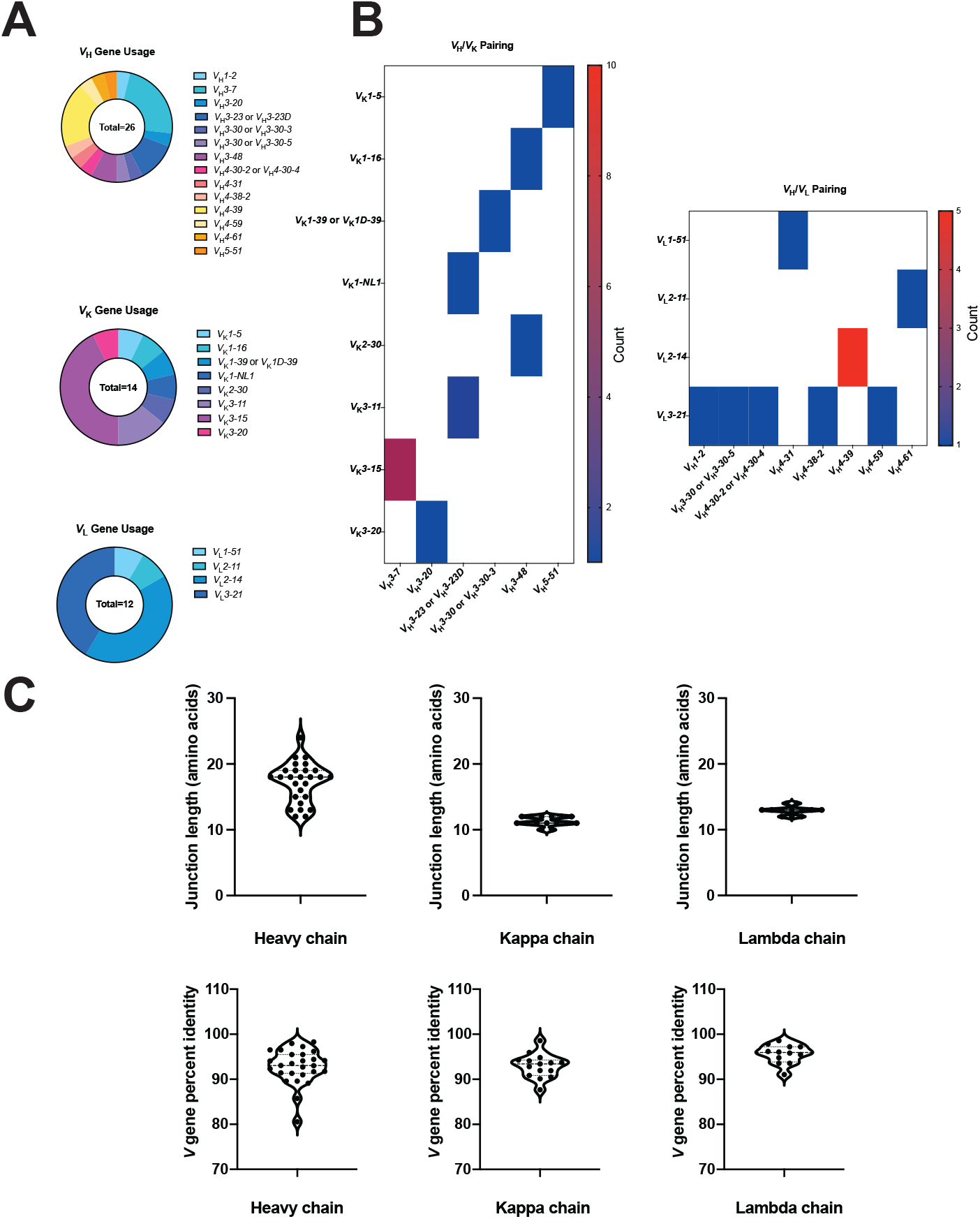
Gene usage and junction lengths of isolated mAbs. (A) The usage of heavy, kappa, and lambda chain genes are shown as a proportion of all respective genes from the panel of isolated mAbs by sequencing the hybridoma line for each clone. The pairing of heavy and light chains is shown in (B), with the number of antibodies corresponding to each pairing shown as a heat map. (C) The amino acid lengths of the junction for the heavy and light chains are shown (top) alongside the percent identity of the *V* gene to the germline sequences (bottom).

### Binding analysis of COBRA HA-specific human mAbs

The majority of isolated mAbs demonstrated high binding to A/California/04/2009 HA protein by ELISA, with an average EC_50_ of 30 ng/mL **(Figure 3A, Figure S1)**. Of these A/California/04/2009 HA protein-reactive mAbs, only a subset demonstrated binding to the P1 and X6 COBRA HA proteins. mAbs P1-02, P1-05, and 163-20 showed reactivity against the P1 COBRA protein, and mAbs CA09-26, CA09-30, CA09-45, P1-02, and P1-05 demonstrated significant binding to the X6 COBRA protein. The limited mAb binding to P1 and X6 COBRA proteins correlated with the lower reactive B cell frequencies to these respective HAs in **Figure 1.** No binding was observed for any mAb to the H3 subtype HA HK14 or to an irrelevant antigen control **(Figure S1)**. Recently, a next-generation H1 COBRA protein, Y2, was generated, which shares 97.5% sequence identity with A/California/04/2009 HA **(Figure 4)** (37). All mAbs had similar EC_50_ values and reactivity to the Y2 COBRA protein compared to A/California/04/2009 HA protein **(Figure 3A, Figure S1)**. We also determined if mAbs target the stem region by utilizing a chimeric HA protein bearing a H6 HA head and a H1 HA stem (cH6/1) **(Figure 3A)**. mAb P1-05 bound to the chimeric protein with high affinity, suggesting this mAb target the stem region of the H1 HA protein. These results indicate that 2017-2018 QIV-vaccinated subjects developed mAbs with potent binding to the 2009 pandemic-like Y2 COBRA HA protein, and each of these subjects produced at least one mAb with reactivity against one or more homosubtypic COBRA HAs. Based on these data and the B cell screening data, COBRA-reactive B cells constitute a major portion of the human B cell response to influenza vaccination, and COBRA HA antigens can likely recall B cells targeting both the head and stem regions.

**Figure 3.**
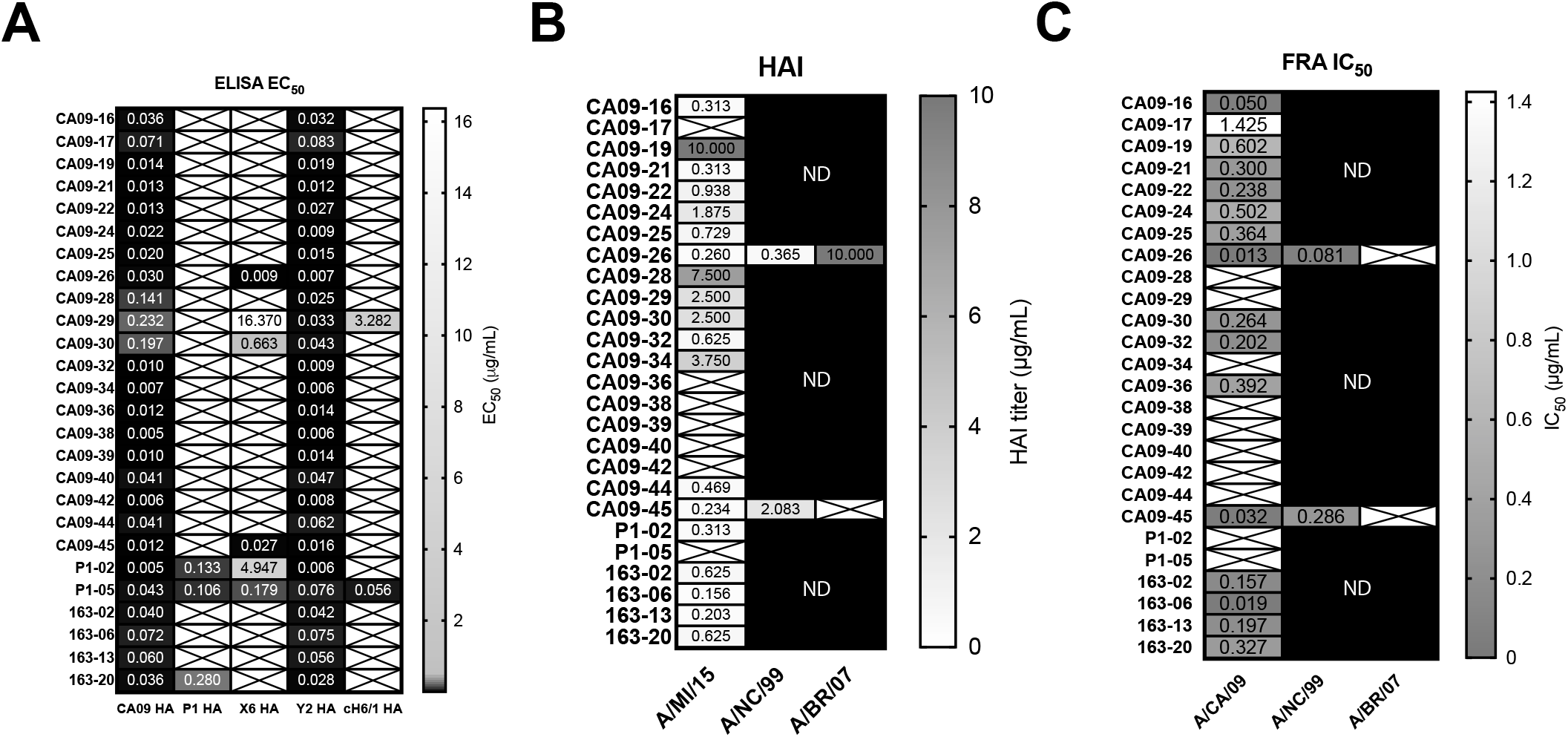
Reactivity of mAbs isolated from 2017-2018 QIV-vaccinated subjects. (A) Half-maximal effective concentrations (EC_50s_) are represented for each mAb. ELISAs were completed with each mAb serially diluted three-fold. Shown are the EC_50_ values against CA09 HA (CA09), P1 COBRA HA, X6 COBRA HA, Y2 COBRA HA, and cH6/1 HA proteins. For the EC_50_ heat map, boxes with an X indicate the signal at 20 µg/mL did not reach 1.5, or the calculated EC_50_ was outside the tested concentration range due to an overall low signal. In (B) boxes with an X indicate no HAI activity was observed at 10 µg/mL. In (C), boxes with an X indicate less than 50% neutralization at the highest concentration tested or the calculated IC_50_ was outside the tested concentration range due to overall low neutralization activity.

**Figure 4.**
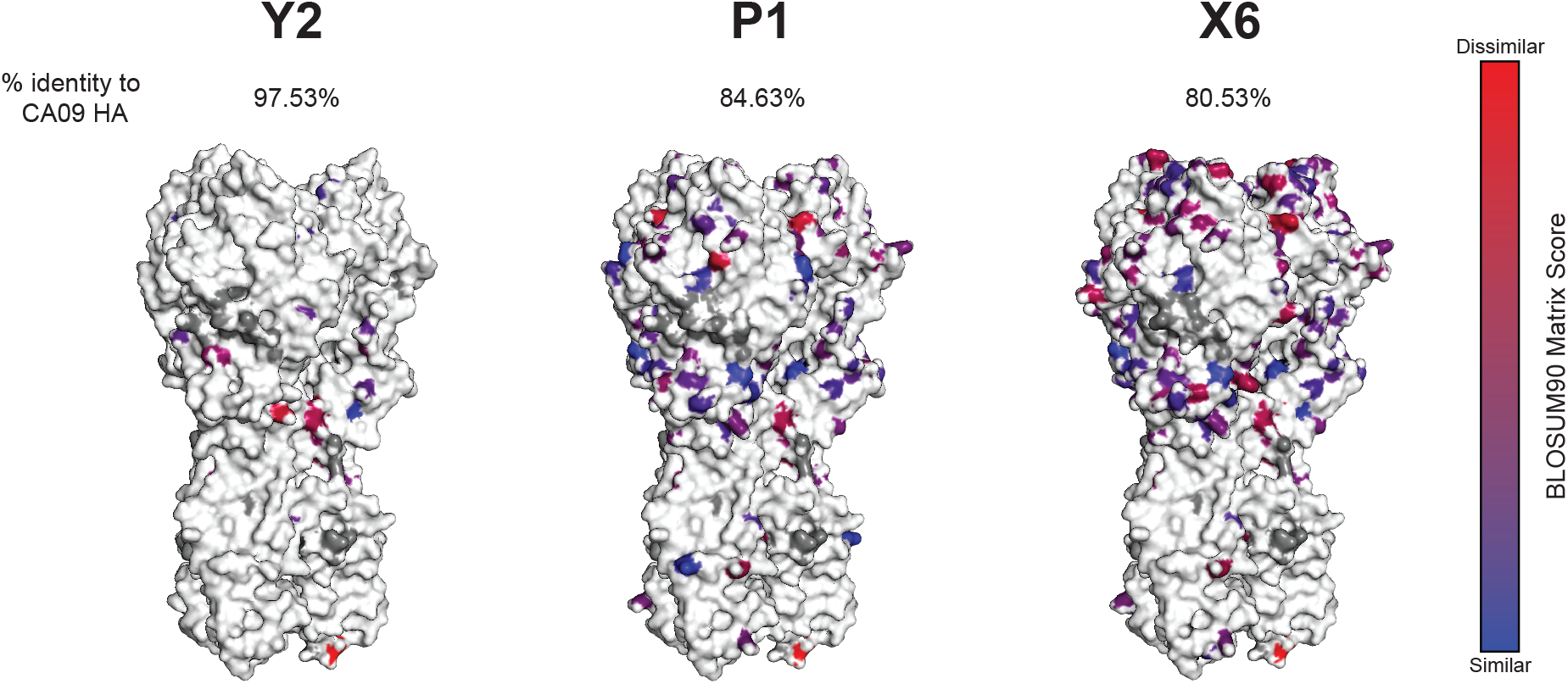
Models of H1 COBRAs used in this study. The models of the H1 subtype COBRAs used are shown alongside the percent identity to the A/California/04/2009 HA. Substitution mutations are indicated in colors corresponding to the BLOSUM90 matrix score, a measure of the likelihood of a given amino acid mutation. White residues indicate an identical amino acid as the A/California/04/2009 HA, blue a substitution with a highly similar amino acid, and red a substitution with a highly dissimilar amino acid. Models were generated using SWISS-MODEL.

### Functional analysis of COBRA HA-specific mAbs

To characterize the functional activities of the isolated mAbs, HAI and neutralizing activities were assessed **(Figure 3B, 3C, S2)**. The majority of mAbs showed HAI activity against the pandemic-like A/Michigan/45/2015 virus **(Figure 3B)**. These data are consistent with the fact that most mAbs bound the head domain of A/California/04/2009 HA **(Figure 3A)**. Of those mAbs with the highest HAI activity of the panel against the recent pandemic-like A/Michigan/45/2015 virus, CA09-26 and CA09-45 were tested for HAI against two pre-pandemic H1 viruses as these mAbs bind the X6 HA COBRA, which incorporates pre-pandemic sequences, and both target the HA protein head domain. CA09-26 had HAI activity against A/New Caledonia/20/1999 and A/Brisbane/59/2007 viruses, while CA09-45 had HAI activity against A/New Caledonia/20/1999, and no activity against A/Brisbane/59/2007 **(Figure 3B)**. We next assessed neutralizing activity against A/California/07/2009 **(Figure 3C**, **S2)**. Approximately 60% of mAbs (16/26 mAbs) neutralized the pandemic A/California/07/2009 virus (A/CA/09). Notably, mAbs CA09-26 and CA09-45 were among the most potent mAbs in the panel with half-maximal inhibitory concentrations (IC_50_s) of 0.014 μg/mL and 0.032 μg/mL, respectively. These two mAbs were also tested for neutralizing activity against A/New Caledonia/20/1999 and A/Brisbane/59/2007 **(Figure 3C)**. They had IC_50_ values of 0.081 μg/mL and 0.286 μg/mL, respectively, against A/New Caledonia/20/1999, indicating potent neutralization activity **(Figure S2)**. However, these two mAbs did not demonstrate neutralizing activity against the A/Brisbane/59/2007 virus, in accordance with the observation that little to no HAI activity was observed for the same strain.

Neutralization-independent, Fc-dependent activities are an important aspect of anti-influenza antibodies that bind both the head and stem domains (38–40). Stem-binding antibodies elicited by P1 HA vaccination also demonstrate Fc activity by inducing cellular cytotoxicity (16). To determine the extent of one such Fc effector function, antibody-dependent phagocytosis (ADP) activity was measured by assessing the capacity for the monocytic THP-1 cell line to phagocytose Y2 COBRA HA-coated beads through mAb binding **(Figure 5)**. The entire mAb panel demonstrated ADP activity relative to the negative mAb control. These included both neutralization/HAI-positive mAbs as well as mAbs that did not demonstrate significant HAI or neutralization activity.

**Figure 5.**
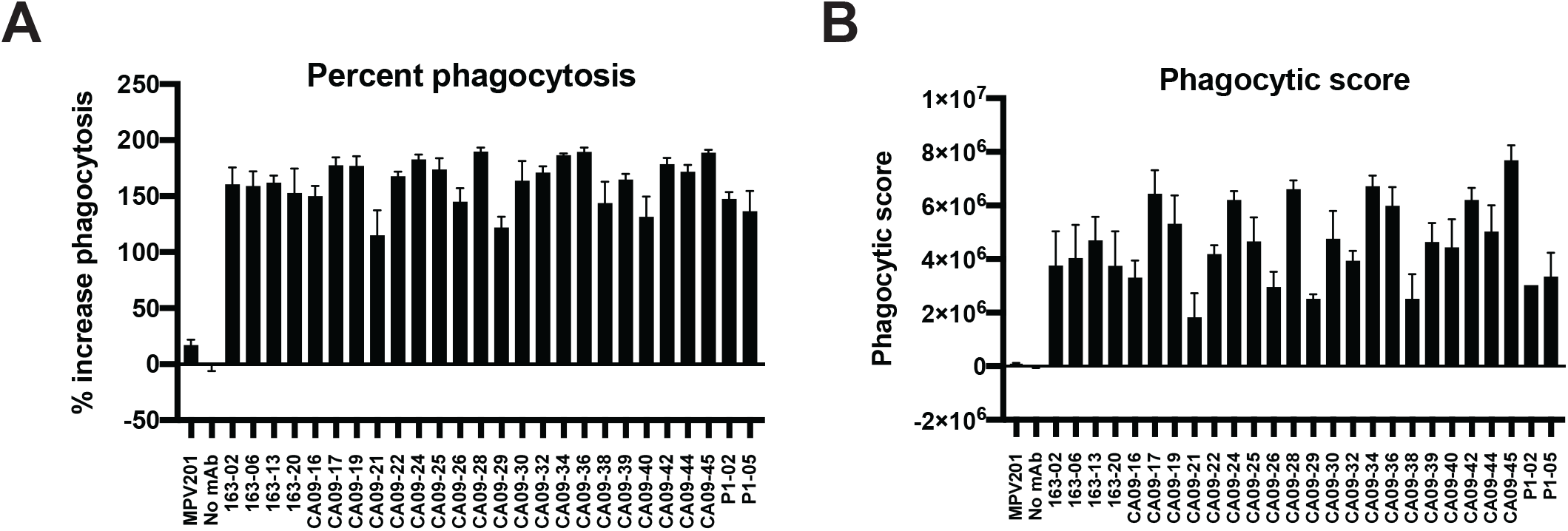
ADP activity of isolated mAbs. The percent phagocytosis of Y2-coated beads by THP-1 cells in the presence of each mAb was assessed using a flow cytometric assay. The relative percent increase of phagocytic cells for each mAb relative to the no mAb control (A), in addition to the phagocytic score (B), are shown. Bars represent the average of three replicates, while errors bars are the standard deviation.

### Multiple distinct epitopes on the A/California/04/2009 HA are bound by isolated mAbs

To determine the epitopes bound by the panel of 27 mAbs isolated from these vaccinated subjects, biolayer interferometry-based epitope binning was utilized as previously described (24, 41). Biosensors were loaded with A/California/04/2009 HA protein, associated with one mAb, and then exposed to a second mAb to determine mAb competition **(Figure 6)**. Control mAbs Ab6649, 5J8, and CR6261 were utilized to determine the relative locations of each epitope. Ab6649 binds the lateral patch, proximal to the Sa antigenic site; MAb 5J8 binds the receptor-binding site (RBS), comprising antigenic sites Sb and Ca2; and mAb CR6261 binds a conserved portion of the stem region found for all group 1 viruses **(Figure 6A)**. Five distinct epitopes on A/California/04/2009 HA protein were distinguished **(Figure 6B)**. Of the epitopes on the globular head domain, two known major epitopes, termed epitope 1 and epitope 3, corresponding to those of Ab6649 and 5J8, respectively, were identified. The position of one predominantly bound epitope, epitope 2, could not be identified by epitope binning with the control mAbs used. Three other epitopes, characterized only by the competition of a single mAb to itself, were epitopes 4 and 5, which correspond to mAbs CA09-38 and P1-05, respectively. No mAbs competed with CR6261, indicating that although mAb P1-05 targets the stem, as evidenced by binding to the cH6/1 protein **(Figure 3, S1),** this mAb targets a different epitope on the stem of the H1 HA protein. Overall, these data suggest that the epitopes bound by mAbs from vaccinated subjects are comprised, in part, of conserved sites on the head domain, such as those involving the RBS and the lateral patch, in addition to portions of the stem.

**Figure 6.**
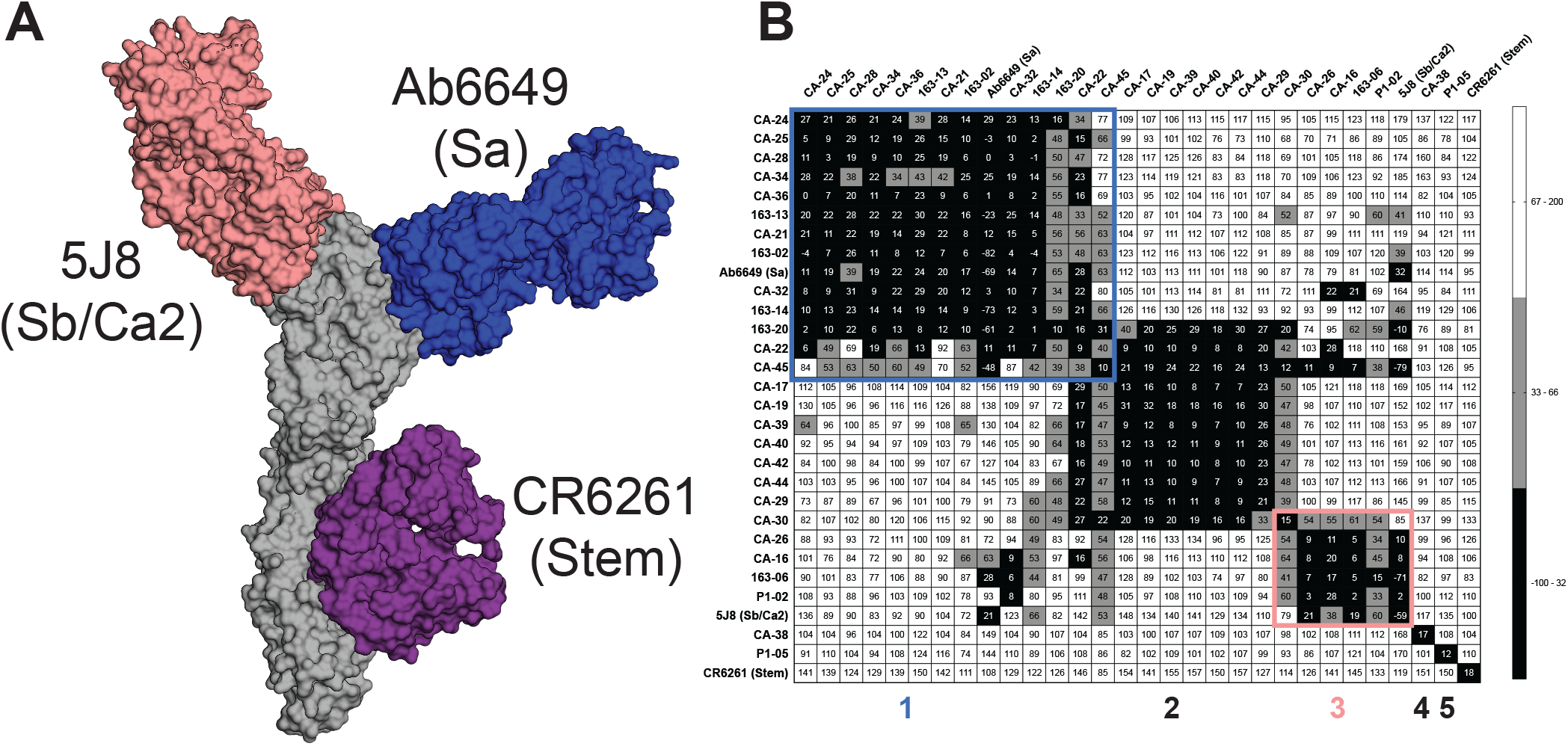
Epitope binning identifies five epitopes from human antibodies isolated at 21 days post-vaccination. (A) Model of A/California/04/2009 HA in complex with three control antibodies used for epitope binning. 5J8 and Ab6649 bind the head domain at the conserved RBS and lateral patch epitopes, respectively. CR6261 binds the stem domain at a site conserved for group 1 viruses. (B) Epitope binning was performed against A/California/04/2009 HA. Competition was measured as the percentage of the response from the association of the second antibody (horizontal axis) in the presence of the first antibody (vertical axis) as compared to the second antibody alone. Black indicates complete competition, gray moderate competition, and white no competition. Identified epitopes that have been previously characterized are outlined in blue (for the lateral patch) and pink (for the RBS). The antigenic sites of the epitopes of control mAbs Ab6649 and 5J8 are shown in parentheses.

Epitope 2 encompassed the binding regions of several mAbs, yet did not compete significantly with any of the control mAbs used. To further elucidate its position on the A/California/04/2009 HA protein, escape mutants of the A/California/07/2009 virus were generated **(Figure 7)**. Following sequencing of the HA gene of these escape mutant viruses, a number of cell adaptation mutations (G165E and S193P) were identified, in addition to escape mutations. The positions of these escape mutations correlated with the expected positions based on the BLI-based epitope binning assay. I176F, D178N, and L201I mutations were found in viruses passaged with the 163-13 mAb that binds epitope 1. 163-13 competed with the Ab6649 antibody, which makes contacts with the Sa antigenic site, and also appears to sterically hinder the Ca2 antigenic site. These mutations correspond to the Ca2 site, and therefore corroborate the epitope binning results for epitope 1. Two other mAbs binding to epitope 1, 163-20 and CA09-22, generated a mutation proximal to Sa, K129N. Interestingly, CA09-30, an epitope 2-binding mAb, also led to the generation of the K129N mutation, also found in escape mutants to epitope 1. This mutation (called K136N with H1 numbering) has previously been reported to introduce a glycosylation site, which may lead to shielding of the antibody epitope (42). It is possible that this epitope is present between epitopes 1 and 3 on the HA head domain, and that this K129N mutation may be a method of virus escape from mAbs that bind in different orientations within these distinct epitopes. No escape mutations were found for any mAbs in epitope 3. Epitopes 4 and 5, which are defined by mAbs CA09-38 and P1-05, respectively, did not generate any escape mutations.

**Figure 7.**
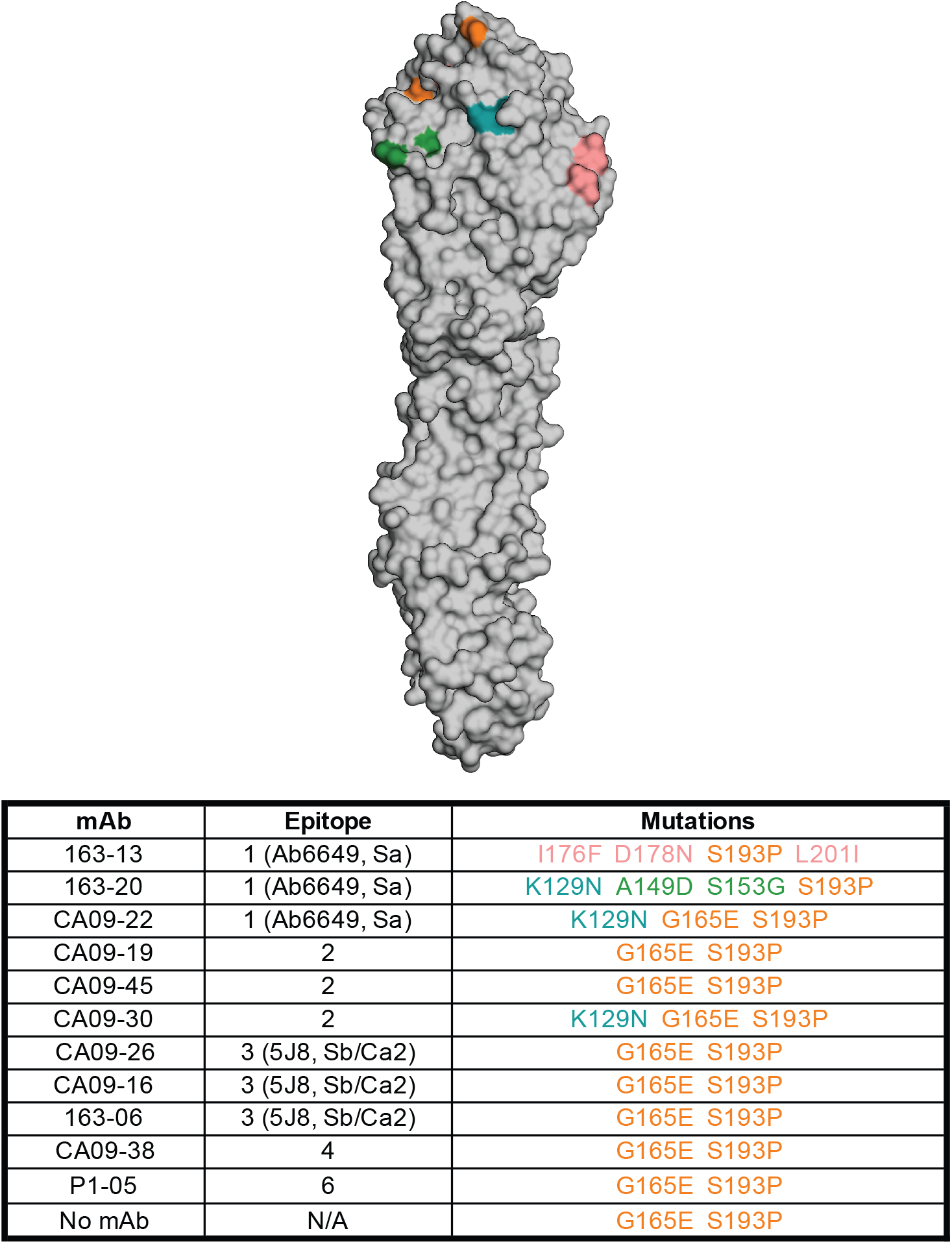
Escape mutants generated from serial passage with H1-reactive mAbs identify their epitopes. The mutations generated through serial passage of the A/California/07/2009 virus in the presence of the indicated mAbs are mapped onto the structure of the H1 HA (PDB ID 5GJS). The mutations are reported in the bottom table for each mAb and are colored as on the HA. The numbering scheme for the PDB 5GJS structure is used.

Epitope 5 is located on the stem of the H1 HA protein yet does not overlap with the conserved stem epitope of mAb CR6261 **(Figure 3A, 6)**. To determine the epitope of mAb P1-05, we generated a complex of Y2 HA bound to P1-05 Fab fragments and evaluated its structure by negative-stain electron microscopy (EM) **(Figure 8A, 8B)**. The 2D class averages revealed that P1-05 binds to the base of the HA stem in an upward angle **(Figure 8B)**. We also observed that insertion of residues between the Y2 C-terminus and the Foldon trimerization domain disrupted mAb P1-05 binding, potentially due to trimer splaying and disruption of this membrane proximal epitope **(Figure 8C)**. Recently, a similar class of mAbs targeting this region on HA, termed the “anchor” epitope, was discovered, and such mAbs protect against H1N1 infection in mice (43, 44). Anchor mAbs do not compete with known stem mAbs and utilize *V_K_3-11* or *V_K_3-15* kappa V genes that encode a germline encoded NWP motif in the CDR3 region (44). The restricted light chain usage can pair with *V*_H_*3-23*, *V*_H_*3-30/V*_H_*3-30-3*, and *V*_H_*3-48* V genes. mAb P1-05 utilizes *V_K_3-11* paired with *V*_H_*3-23*. Furthermore, it was also recently reported that binding of anchor mAbs is disrupted by the use of a GCN4 trimerization domain (44), which has different spacing than the Foldon domain, which matches our data with the disruption of binding observed in **Figure 8C.** These observations are critical for subunit HA protein vaccine development as they indicate the importance of antigen design, stability, and the incorporation of mAb binding affinity studies to ensure that important epitopes are properly displayed on candidate vaccine antigens.

**Figure 8.**
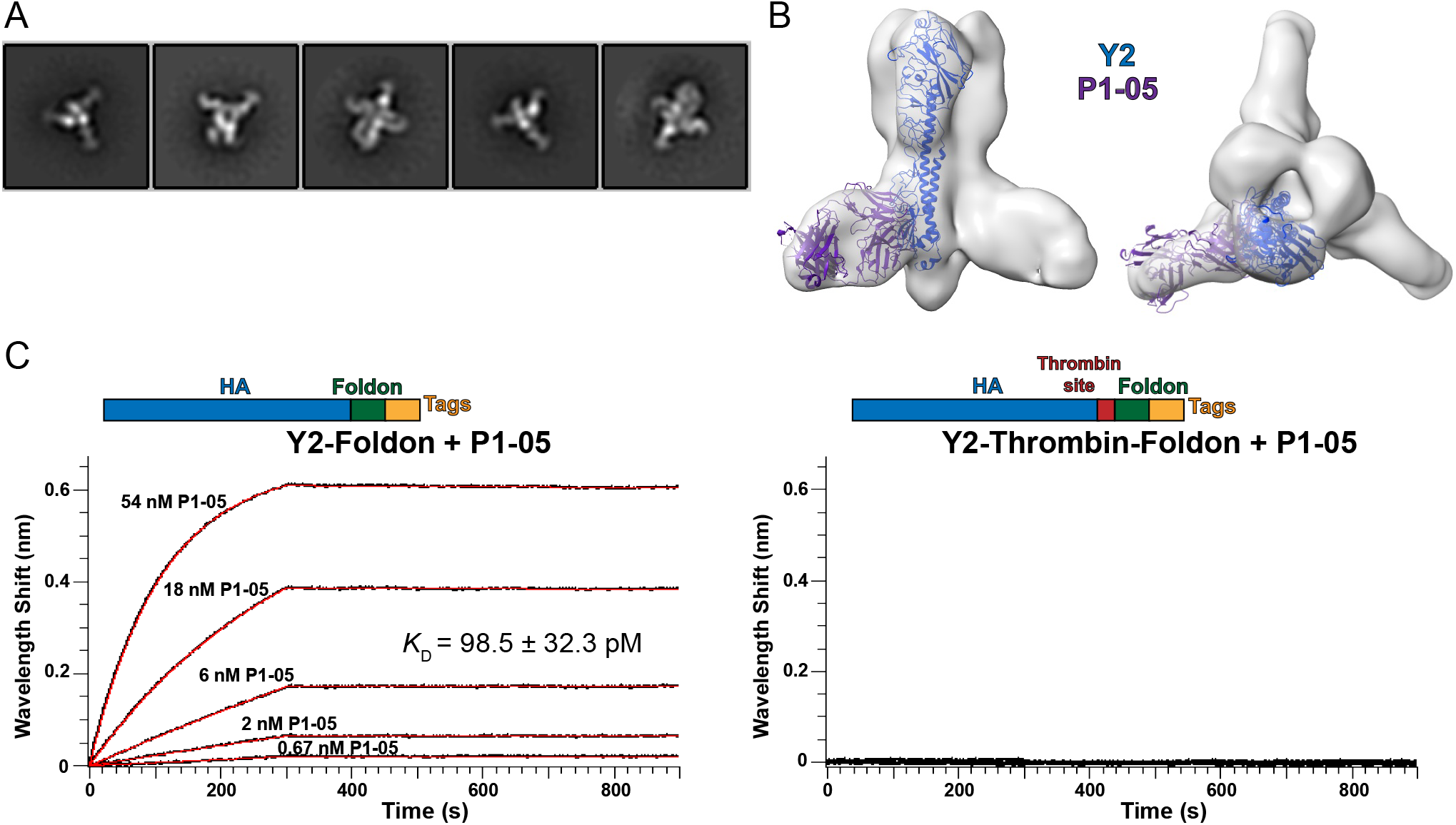
Structural characterization of P1-05 binding to Y2 COBRA. (A) 2D class averages and (B) 3D reconstruction of the Y2+P1-05 complex. (C) Comparison of P1-05 binding with Y2 in the presence or absence of a thrombin cleavage site by biolayer interferometry. Representative runs are shown. The *K*_D_ represents the mean ± standard deviation of three independent experiments.

### Repertoire analysis of Y2-specific B cells

To further probe the repertoire of pre-existing COBRA HA-specific B cells and to determine the prevalence of each mAb in the public antibody domain, we conducted a single-cell RNA sequencing experiment using B cells from a single subject vaccinated with the 2019-2020 seasonal influenza vaccine. Approximately 3000 CD19^+^IgM^-^IgD^-^ B cells positive for the Y2 COBRA HA were sorted and subjected to 10X barcoding **(Figure 9, S3)**. Prior to loading onto the 10X controller, sorted Y2-specific cells were supplemented with the CA09-26 hybridoma clone as a loading control. 69 unique paired heavy and light chains were obtained following data demultiplexing and analysis compared to the human genome database. Similar to the mAb sequencing, the *V*_H_*1* and *V*_H_*4* gene families were highly prevalent in the B cell repertoire. In particular, *V*_H_*4-39* and *V*_H_*4-59* were prevalent in both mAb sequencing and B cell sequencing results. We also identified several additional mAbs utilizing the *V*_H_*3-23* gene, with one in particular (clone 70) having an NWP motif in a paired *V_K_3-15* light chain, consistent with an additional anchor-like mAb (44) **(Table S1)**. A fraction of *V*_H_*1-69* genes were also identified, which is utilized by mAbs targeting the stalk epitope. Hence, we observed a relatively diverse repertoire of binding antibodies including both central stalk and anchor targeting mAbs.

**Figure 9.**
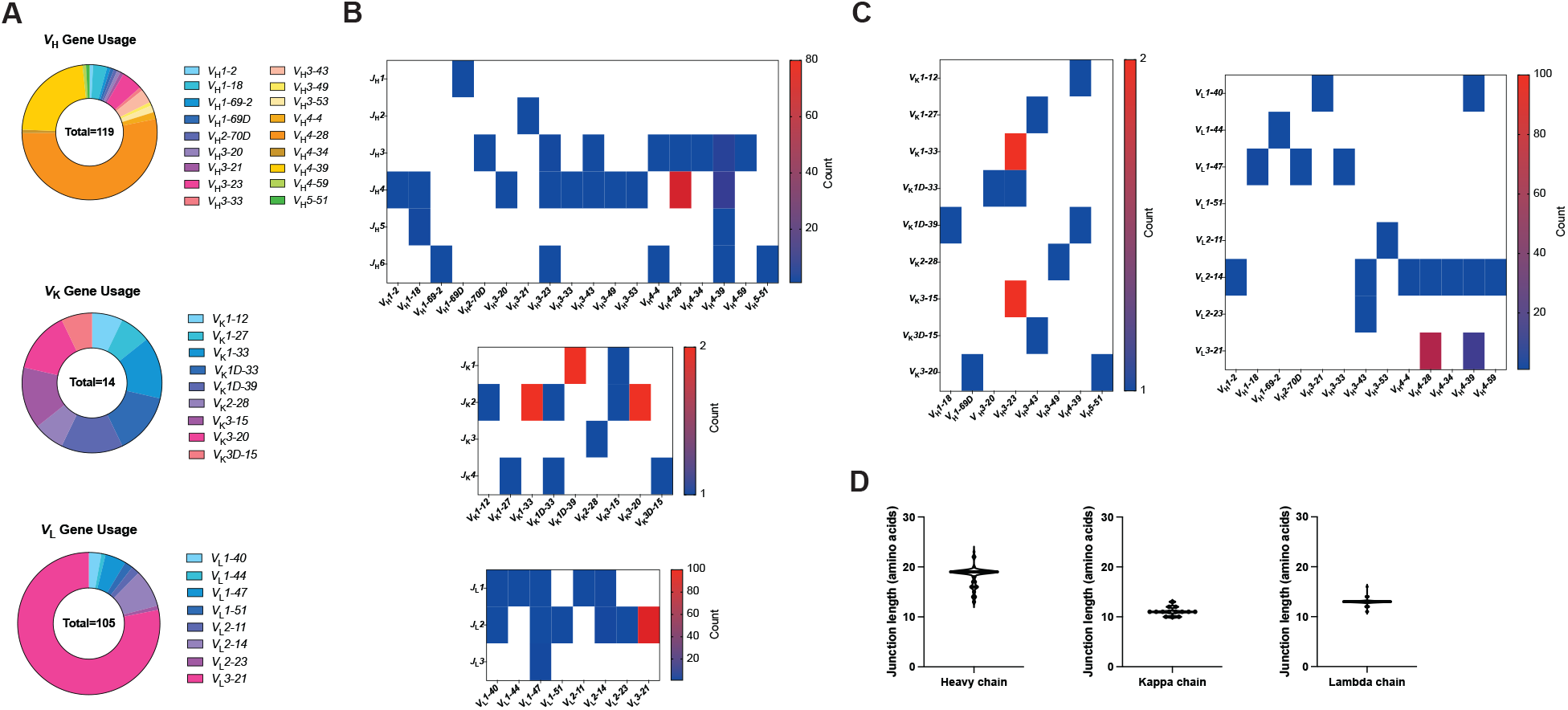
Sequence characteristics of Y2 COBRA-specific B cells from a human subject receiving the 2019-2020 seasonal vaccine. (A) The usage of heavy, kappa, and lambda chain genes are shown as a proportion of all respective genes for all B cells with paired heavy and light chains. (B) The pairing of *V* and *J* genes are shown, with the number of B cells contributing to each pairing for each chain, as heat maps. (C) The pairing of heavy and light chain *V* genes is shown for heavy-kappa chain pairings (left) and heavy-lambda chain pairings (right) as heat maps. (D) The amino acid lengths of the junctions for the heavy and light chains are shown.

## Discussion

H1 COBRA HA antigens have been successful at broadening the antibody response compared to wild-type HA sequences in naïve and pre-immune mouse and ferret models of influenza infection (22, 37). However, pre-existing immunity to influenza in humans remains a major challenge to overcome due to repeated previous exposure to the influenza HA protein during infection and vaccination events. In this study, we sought to determine the extent of the H1 subtype COBRA HA-reactive pre-existing B cell repertoire in human subjects to predict recall responses as COBRA HA antigens move toward clinical trials. At the oligoclonal B cell level, pre-existing B cell responses were observed for P1 and X6 COBRA antigens in individuals vaccinated with the 2017-2018 seasonal influenza vaccine, which incorporated the pandemic-like A/Michigan/45/2015 vaccine strain. COBRA HA-reactive B cell responses were lower than those observed for A/California/04/2009 HA protein, likely due to loss of strain-specific variable head epitopes and incorporation of seasonal pre-pandemic and swine HA sequences in the X6 and P1 antigens, respectively. mAbs isolated against A/California/04/2009 HA protein utilized a diverse gene repertoire and only a small subset reacted with P1 and X6 COBRA antigens. In contrast, the mAb binding profile to the recently described Y2 COBRA HA, which utilizes 2009 pandemic-like H1 sequences from 2014-2016, was similar to that observed for the A/California/04/2009 HA protein. The majority of the mAbs had HAI activity and neutralizing activity against A/Michigan/45/2015 and A/California/07/2009, and two head-binding mAbs that bind the X6 protein, CA09-26 and CA09-45, had HAI activity and neutralizing activity against the pre-pandemic strain A/New Caledonia/20/1999. These data suggest that the COBRA X6 HA-reactive mAbs are mainly endowed with functional activity against both pre-pandemic and pandemic-like H1 viruses. Overall, the amino acid similarity of COBRA HA antigens to A/California/04/2009 HA correlated with high B cell and mAb reactivity. In addition to binding, neutralization, and HAI activity, we also assessed if COBRA HA-reactive mAbs had Fc-mediated functions, namely ADP, and virtually all mAbs were able to induce THP-1 phagocytosis of Y2-coated beads.

Several epitopes on the H1 HA protein have been previously defined (10), and we determined mAb epitopes on the A/California/04/2009 HA protein using biolayer interferometry. The majority of the mAbs targeted three head-binding epitopes on the Sa and Sb/Ca2 sites, and an undefined epitope identified through the generation of escape mutations. CA09-38 did not exhibit HAI or neutralizing activity, nor did it bind the cH6/1 HA, suggesting that this mAb targets an undefined non-neutralizing epitope on the head region. We discovered that P1-05 targets a unique epitope on the H1 HA stem region, and this epitope is similar to the recently described anchor epitope (44). Based on these data, while COBRA antigens were primarily designed to induce broadly reactive antibodies to the head domain, these antigens will likely also recall broadly reactive anchor mAbs in humans in addition to head-based recall and *de novo* antibody responses. Further repertoire analysis in a subject vaccinated with the 2019-2020 seasonal influenza vaccine identified additional anchor-like mAb sequences, as well as similar sequences to our mAbs targeting the head domain from the 2017-2018 season, indicating that the COBRA- reactive B cell population is similar across subjects and influenza vaccine seasons.

These data provide evidence that a pre-immune population with exposure to the seasonal influenza virus vaccine exhibits B cell reactivity towards conserved epitopes present on COBRA HA antigens. As the COBRA HA platform enters clinical trials, it is likely that head-specific and some stem-specific antibodies will be elicited as part of a recall response. Moreover, the antibody epitopes identified in this work overlap in part with those previously identified on the head domain near the RBS and the lateral patch, in addition to those on the stem. These epitopes are the focus of future structural studies, particularly for those mAbs that cross-react with the X6 COBRA HA as well as with the HA stem domain. Our data also exemplify the importance of structural analysis of protein epitopes to ensure epitopes that elicit broadly neutralizing antibodies, such as the anchor epitope, remain intact following design optimization for subunit HA vaccines.

## Data availability

The 3D reconstruction of Y2 + P1-05 was deposited to the Electron Microscopy Data Bank under deposition ID D_1000260135.

---

**Figure S1.**
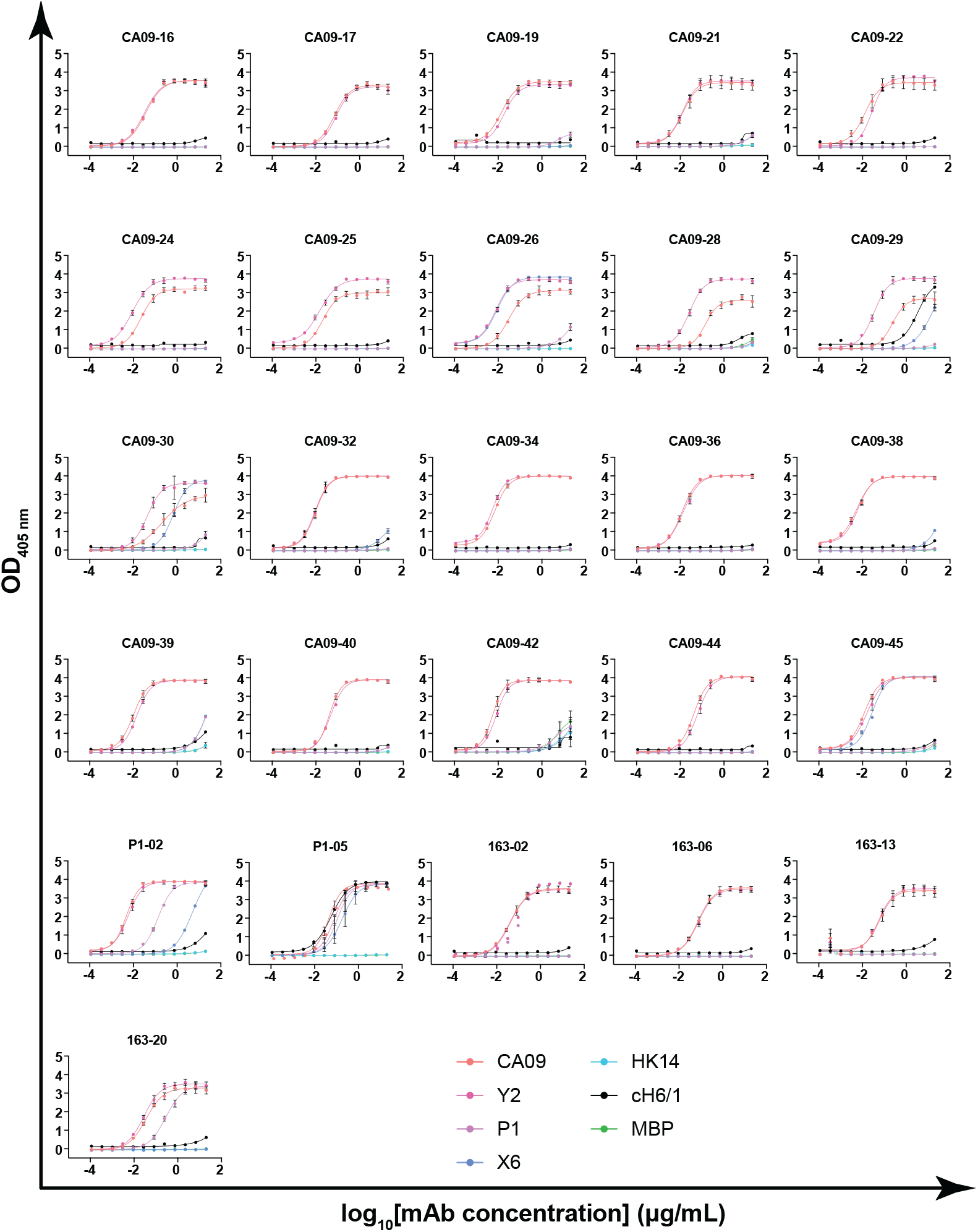
Binding curves for isolated mAbs. The optical density at 405 nm (OD_405 nm_) was measured by ELISA for serial three fold-dilutions of the indicated mAbs from 20 μg/mL. This was assessed for the indicated antigens, A/California/04/2009 HA (CA09), Y2 COBRA HA (Y2), P1 COBRA HA (P1), X6 COBRA HA (X6), A/Hong Kong/4801/2014 HA (HK14), chimeric H6/1 HA (cH6/1), and maltose-binding protein (MBP), a negative control.

**Figure S2.**
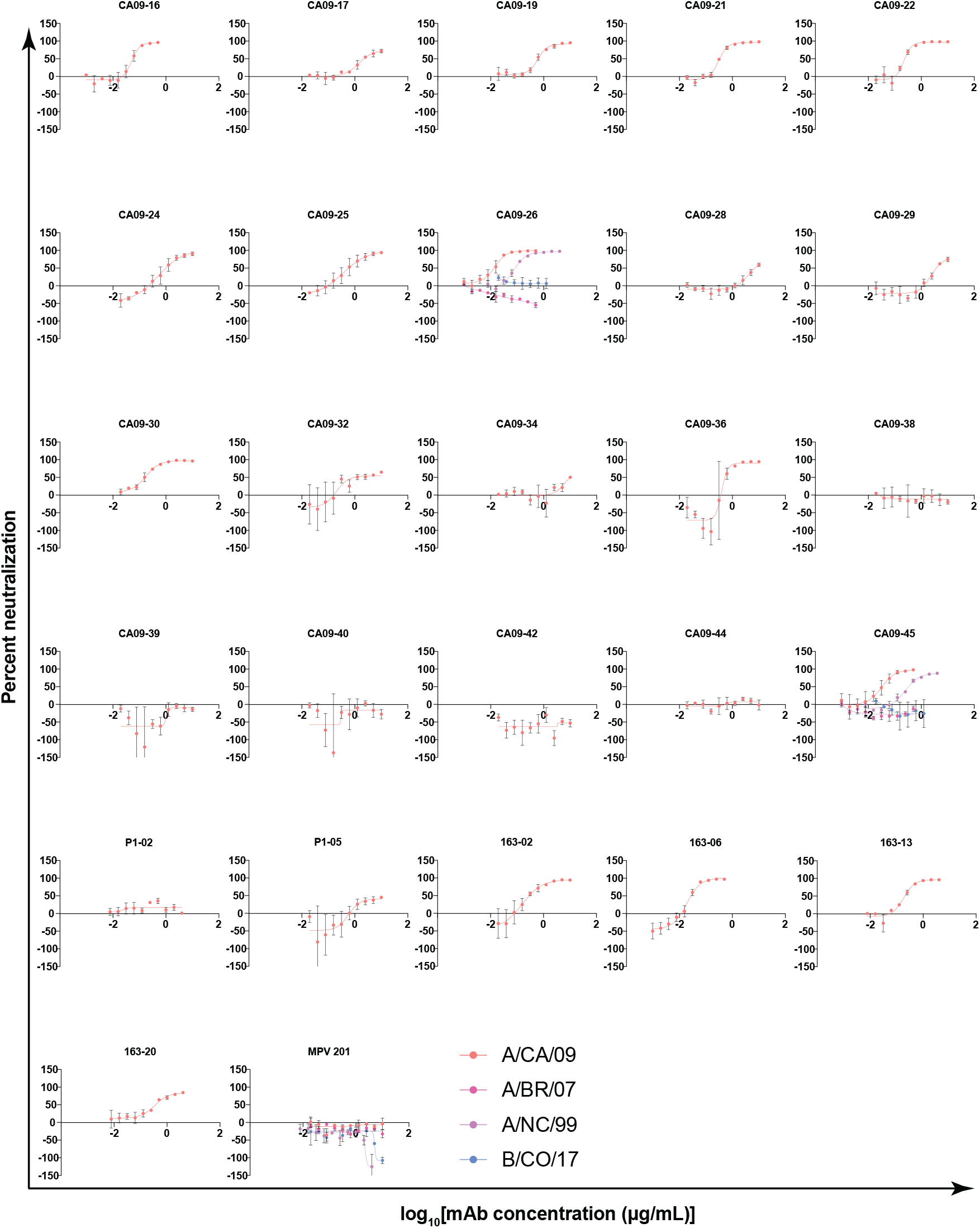
Neutralization activities for isolated mAbs. The percentage neutralization against the indicated H1 subtype influenza viruses was assessed for each antibody. MPV 201 is an irrelevant hMPV-specific antibody.

**Figure S3.**
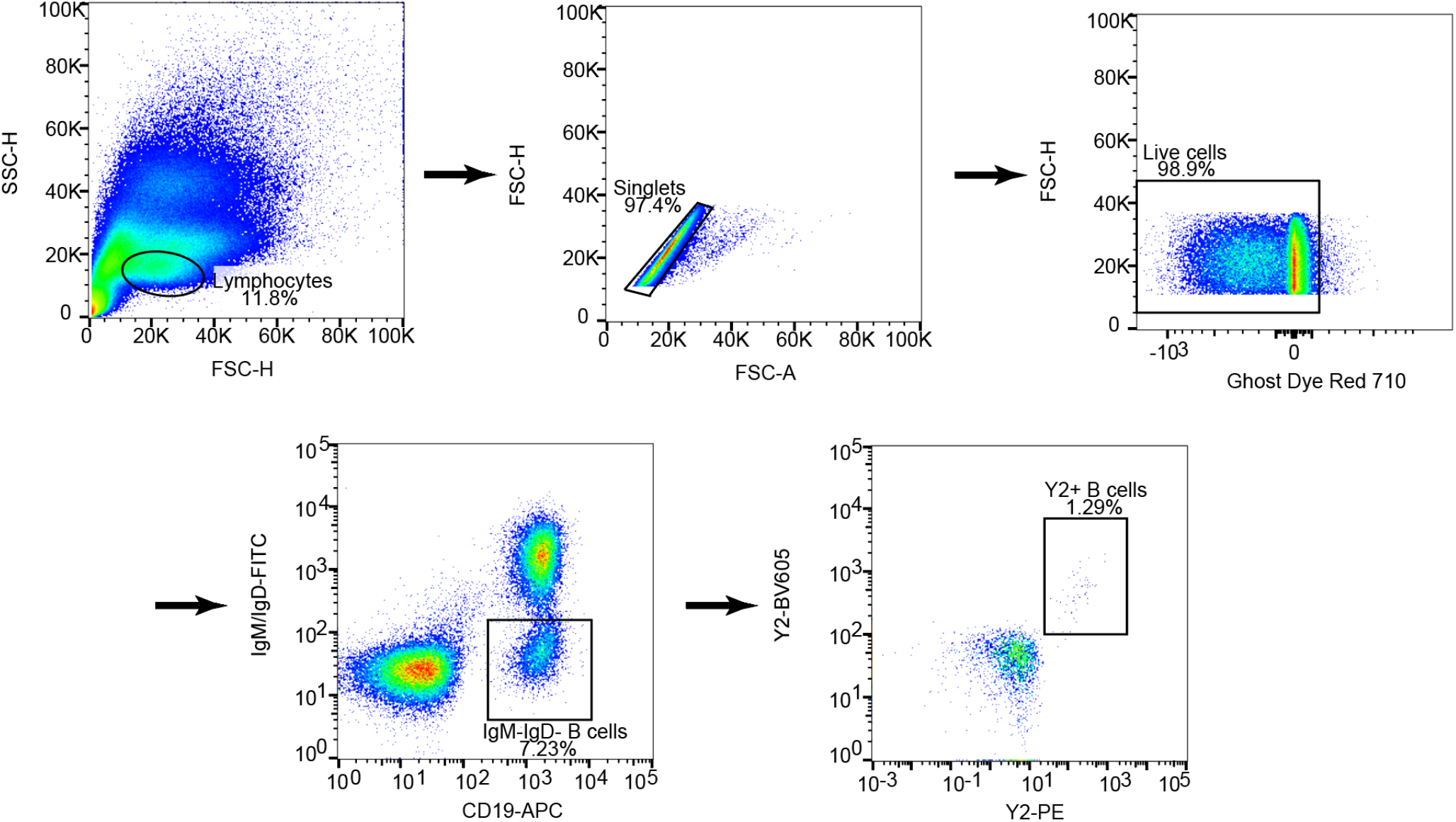
Gating strategy for Y2-specific B cells from PBMCs derived from a subject receiving the 2019-2020 seasonal vaccine. The gating strategy for isolation of Y2-specific B cells from PBMCs is shown. Live, single cells were gated, followed by gating for CD19^+^IgM/IgD^-^ Y2-PE^+^Y2-BV605^+^ B cells.

## Supplementary tables

**Supplementary table S1.**
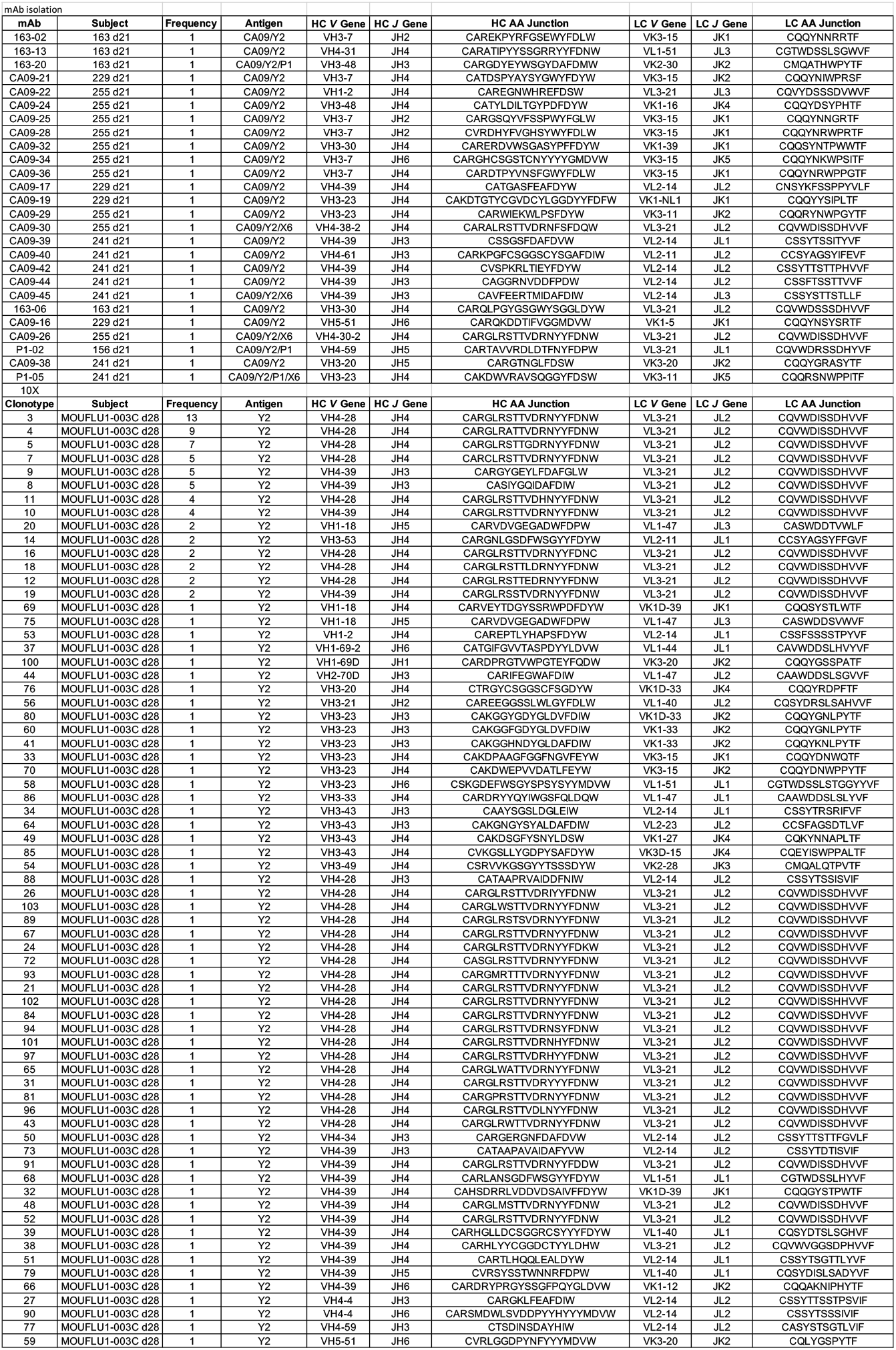
mAb and B cell receptor V(D)J gene usage characteristics. The mAbs elicited by 2017-2018 seasonal vaccination and B cell clonotypes elicited by 2019-2020 seasonal vaccination are shown. The variable (*V*), joining (*J*), and diversity (*D*) genes for the heavy chain (HC) and light chain (LC) are shown, alongside the heavy and light chain isotypes and junction sequences.

